# Modeling the Impact of Seasonality on Mosquito Population Dynamics: Insights for Vector Control Strategies

**DOI:** 10.1101/2024.05.23.595567

**Authors:** Joseph Baafi, Amy Hurford

## Abstract

Mosquitoes are important vectors for the transmission of some major infectious diseases of humans, i.e., malaria, dengue, West Nile Virus and Zika virus. The burden of these diseases is different for different regions, being highest in tropical and subtropical areas, which have high annual rainfall, warm temperatures, and less pronounced seasonality. The life cycle of mosquitoes consists of four distinct stages: eggs, larvae, pupae, and adults. These life stages have different mortality rates and only adults can reproduce. Seasonal weather may affect the population dynamics of mosquitoes, and the relative abundance of different mosquito stages. We developed a stage-structured model that considers laboratory experiments describing how temperature and rainfall affects the reproduction, maturation and survival of different Anopheles mosquito stages, the species that transmits the parasite that causes malaria. We consider seasonal temperature and rainfall patterns and describe the stage-structured population dynamics of the Anopheles mosquito in Ain Mahbel, Algeria, Cape Town, South Africa, Nairobi, Kenya and Kumasi, Ghana. We find that neglecting seasonality leads to significant overestimation or underestimation of mosquito abundance. We find that depending on the region, mosquito abundance: peaks one, two or four times a year, periods of low abundance are predicted to occur for durations ranging from six months (Ain Mahbel) to not at all (Nairobi); and seasonal patterns of relative abundance of stages are substantially different. The region with warmer temperatures and higher rainfall across the year, Kumasi, Ghana, is predicted to have higher mosquito abundance, which is broadly consistent with reported malaria deaths relative to the other countries considered by our study. Our analysis reveals distinct patterns in mosquito abundance across different months and regions. Control strategies often target one specific life stage, for example, applying larvicides to kill mosquito larvae, or spraying insecticides to kill adult mosquitoes. Our findings suggest that differences in seasonal weather affect mosquito stage structure, and that the best approaches to vector control may differ between regions in timing, duration, and efficacy.

## 1 Introduction

Mosquito-borne diseases continue to pose a significant global threat to human health, particularly in tropical and subtropical regions [1, 2, 3]. Malaria, dengue fever, Zika virus, and various other diseases transmitted by vectors impose a significant burden on communities around the globe [4, 5, 6]. Among the vectors responsible for transmitting these diseases, *Anopheles* mosquitoes are of paramount concern due to their role in spreading malaria, which affects millions of people worldwide. Approximately half of the global population is at risk of malaria [1, 7, 8]. Certain parts of the world have successfully eliminated malaria and have been certified malaria-free by WHO (for example, Mauritius in 1973, Morocco in 2010, Croatia in 1973, and Argentina in 2019) [9]. Malaria remains a persistent threat, even in countries with temperate climates, such as the United States, where recent malaria cases have been reported [2]. In 2021, a staggering 247 million cases of malaria were reported in 85 countries, resulting in over 619,000 deaths [1]. A significant majority of these cases occurred in sub-Saharan Africa, with the region accounting for 96% of malaria-related deaths and 95% of malaria cases in 2021 [2]. To control disease spread, it may be necessary to control vector populations in disease-endemic regions. Population dynamics vary regionally and best approaches to vector management, such as mosquitoes, are likely region-specific [10, 11].

Weather conditions, including temperature, precipitation, and humidity, have long been recognized as pivotal determinants influencing mosquito life cycles, population abundance, and the dynamics of disease transmission [3, 12, 13, 14, 15, 16, 17]. The diverse climates and regions across the globe give rise to distinct ecosystems, introducing variability that can impact both mosquito populations and their stage structure. Recent studies have made significant strides by incorporating temperature dependencies into models, accounting for factors such as egg oviposition, mortality, and maturation rates, thus enhancing the predictability of mosquito population dynamics, as demonstrated in studies focusing on *Culex pipiens* [12, 18]. Models that consider temperature dependence have exhibited robust performance across various species within specific regions [12, 19, 20, 21]. The success of these models underscores the importance of investigating the relationship between regional weather variability and mosquito dynamics. Mathematical models that neglect seasonal variations offer simplicity in analysis but may produce inaccurate outcomes and fail to provide adequate insights for determining optimal timing for implementing control measures [11, 22] during different periods of the year.

Various models have been developed to assess the impact of seasonality in environmental conditions on the transmission dynamics of mosquito-borne diseases such as malaria [14, 23] and dengue [17, 20, 24, 25]. Some of these models have delved into mosquito population dynamics under seasonal weather conditions using ordinary differential equation formulations [12, 13, 21, 26, 27] or delay-differential equation systems [28, 29]. However, the majority of existing models are tailored to specific mosquito species within particular geographic contexts. For instance, models have been developed for *Aedes aegypti*, in Brazil [20, 25]; *Anopheles gambiae* in the Sahel region of Africa [30]; *Anopheles arabiensis* in Zambia [31]; *Anopheles arabiensis* in KwaZulu-Natal, South Africa [21] and *Culex* in the Peel Region, Canada [12, 19]). Nonetheless, the generalizability of mosquito population estimates and recommendations for vector control derived from these models may be constrained by variations in regional seasonality and adaptive traits. Despite some studies investigating the impacts of regional seasonality on mosquito population dynamics [32], further investigation in this area is warranted.

For this study, we focus on four distinct regions in Africa: Algeria, South Africa, Kenya, and Ghana. Each region presents a unique climate and ecological context, allowing us to investigate how the interplay of temperature and rainfall influences mosquito population dynamics and stage structure. This study has dual primary objectives. Firstly, we aim to gain a comprehensive understanding of how regional seasonality influences mosquito population dynamics. The integration of temperature and rainfall data into our stage-structured model allows us to assess the impact of these weather variables on mosquito survival, developmental rates, and population growth throughout different seasons. Secondly, we will try to derive insights that can inform effective vector control strategies. By exploring the intricate relationship between environmental conditions and mosquito population dynamics, we can describe periods of peak mosquito abundance. This knowledge enables us to strategically implement interventions during these critical time frames.

The insights obtained from this study hold significance for public health authorities and researchers, guiding the development of context-specific strategies to mitigate mosquito-borne diseases. Potential strategies may include optimizing the timing of insecticide applications, implementing targeted larval source reduction measures, and enhancing surveillance systems to quickly detect and respond to periods of elevated mosquito abundance. This research aims to contribute actionable knowledge, paving the way for more targeted and efficient mosquito control interventions.

The paper is organized as follows. In Section 2, we formulate the model and discuss the data used to find parameter estimates and parameter functions. Section 3 contains numerical simulations and results of the model discussed in section 2.1 using the temperature and rainfall data of four African regions. In Section 4, we discuss the results and their implications and suggest possible future directions. The concluding remarks are presented in Section 5, summarizing key findings and contributions.

## 2 Methods

### 2.1 Model formulation

Mosquitoes progress through four distinct life stages: egg, larva, pupa, and adult. The first three stages occur in water, while adult mosquitoes inhabit land and air [33, 34, 35]. The process begins with egg hatching, leading to the emergence of the first larval instar. Subsequently, the larva undergoes three additional molts, transitioning to the second, third, and fourth larval instars. After completing these stages, pupation occurs, giving rise to the pupa. Finally, the pupa undergoes metamorphosis, leading to the emergence of adult mosquitoes, both male and female. For successful reproduction, mating between male and female mosquitoes is necessary. Copulation results in the fertilization of the female, followed by the oviposition phase, where the adult female lays her eggs after obtaining sufficient blood meal [29, 36, 37].

We formulate and analyze a stage-structured ordinary differential equation model that reflects details of the mosquito life cycle (Figure 1) and considers a seasonal environment (temperature- and rainfall-dependent). The stage structuring corresponds to the three main immature mosquito life stages: egg, *E*(*t*); larva, *L*(*t*); pupa, *P* (*t*), and the adult female mosquitoes, *A*_*M*_ (*t*). We incorporate stage-specific life history rates and processes to describe the rate of transitions between the life stages [28]. We split the adult stage into three sub-stages: adult seeking host, *A*_*h*_(*t*), adult at rest, *A*_*r*_(*t*) and ovipositing adult, *A*_*o*_(*t*) so that *A*_*M*_ (*t*) = *A*_*h*_(*t*) + *A*_*r*_(*t*) + *A*_*o*_(*t*). The model does not explicitly account for adult males but assumes their abundance to be equal to that of adult females. The model is given by the following deterministic system of nonlinear ordinary differential equations (ODE).

**Figure 1:**
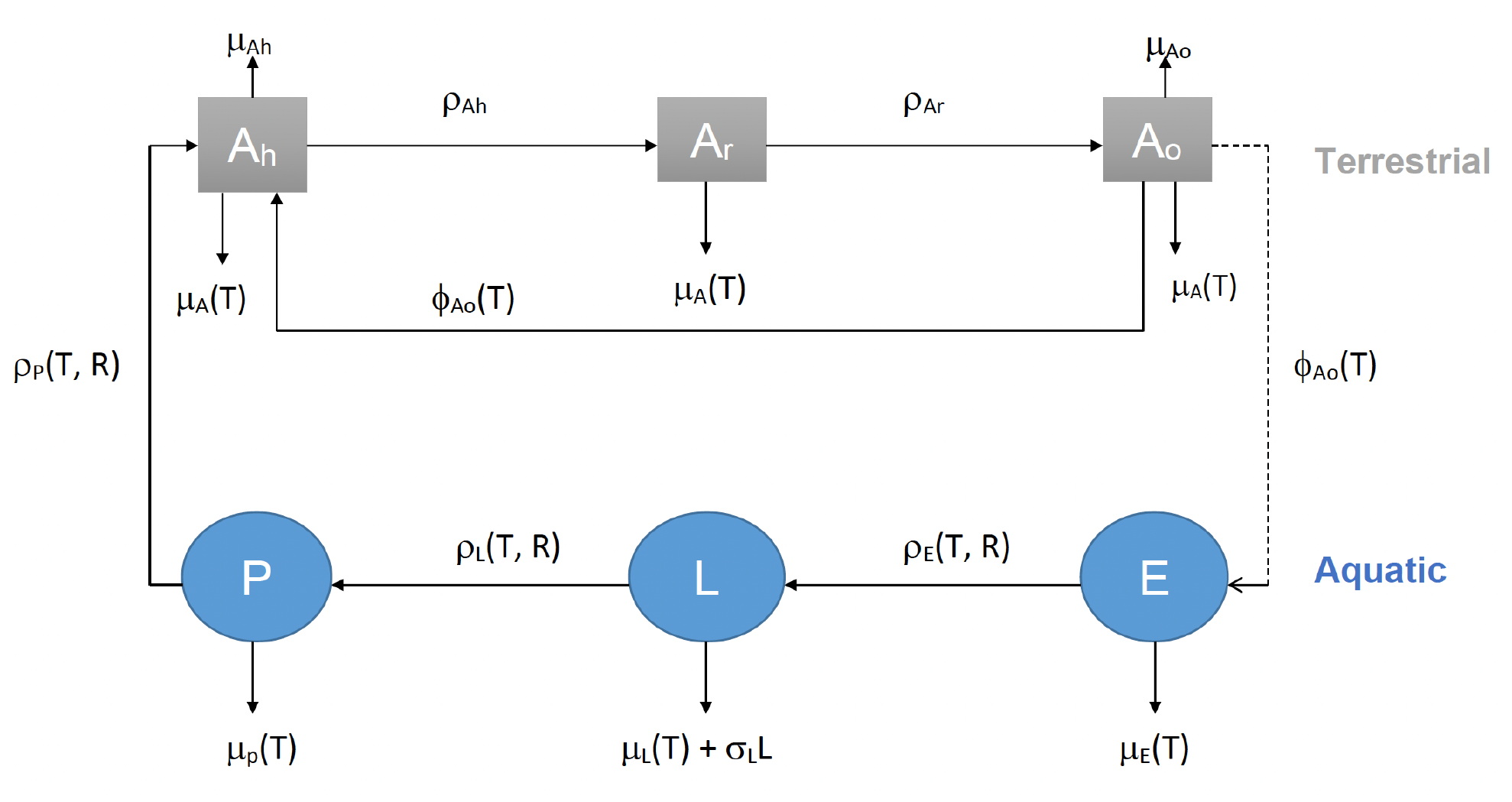
Flow diagram illustrating the model setup (see Equation 2.1), depicting the influence of temperature and rainfall on life history parameters. Density-dependent mortality (*δ*_*L*_) occurs exclusively during the larval stage.

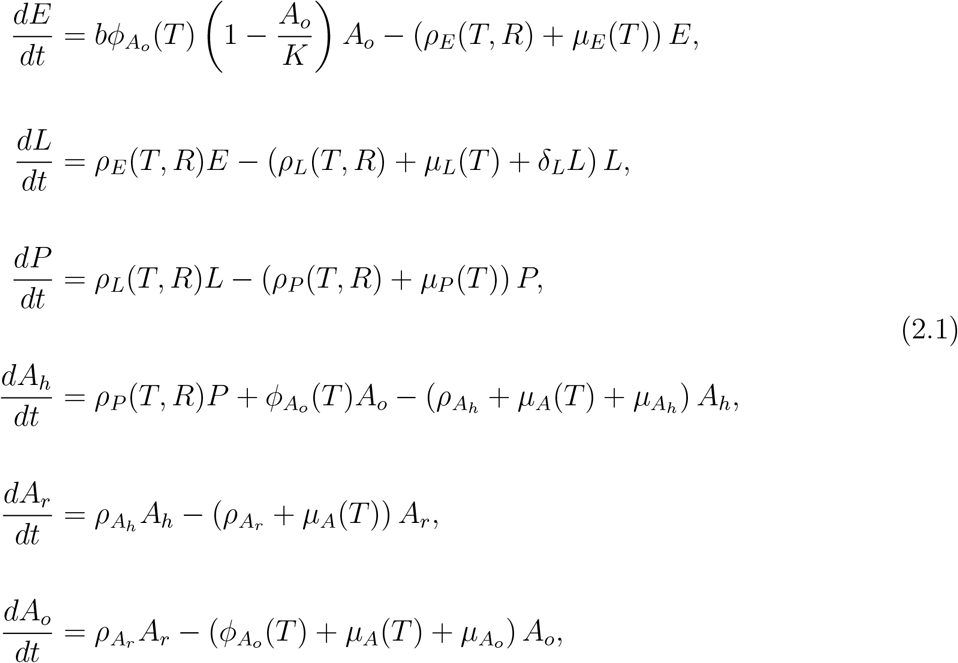

with initial conditions *E*(0), *L*(0), *P* (0), *A*_*h*_(0), *A*_*r*_(0) and *A*_*o*_(0). In system 2.1, *T* = *T* (*t*) and *R* = *R*(*t*) represent temperature and rainfall at time *t*, respectively. We assume that juvenile and adult mosquitoes experience the same temperature (that is, ambient and water temperatures are equivalent near the surface of the water) [12, 28]. Typically, sinusoidal functions, such as

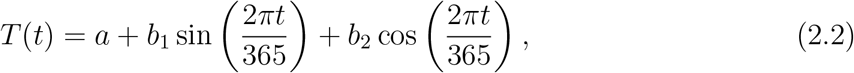

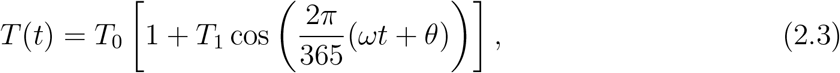

are used to represent regional temperature fluctuations. Similar functions are used to account for rainfall variability (See figure 2). We assume that the functions *T* (*t*) are non-negative, continuous, and bounded periodic functions. Also, the parameters 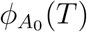, *ρ*_*E*_(*T, R*), *ρ*_*L*_(*T, R*), *ρ*_*P*_ (*T, R*), *µ*_*E*_(*T*), *µ*_*L*_(*T*), *µ*_*P*_ (*T*) and *µ*_*A*_(*T*) are non-negative, periodic, continuous and bounded functions defined on [0, ∞).

**Figure 2:**
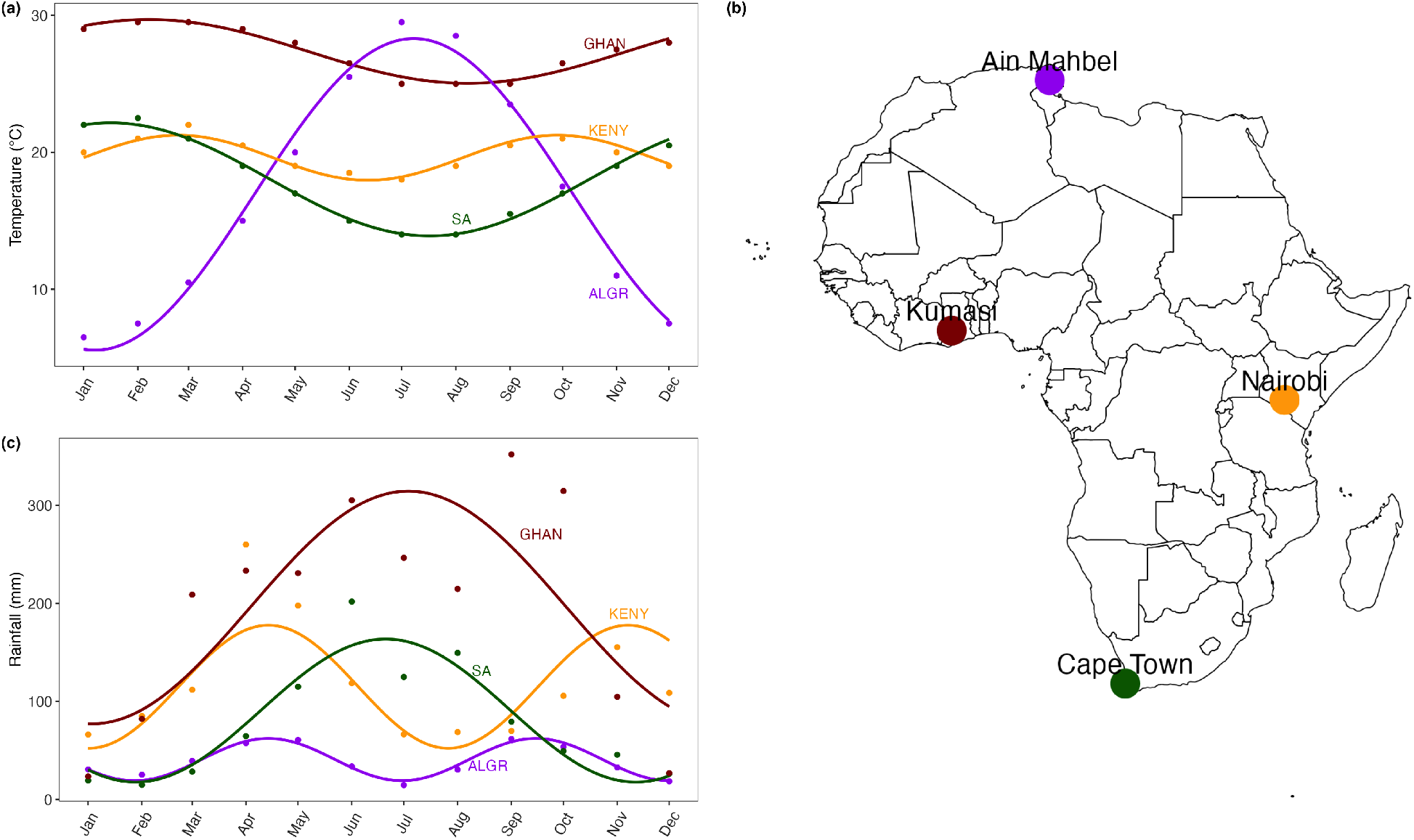
Weather data for the four African regions and their corresponding locations. Panels (a) and (c) depict the average ambient temperature and rainfall patterns with each month’s average taken across twelve years (2009-2020). Panel (b) shows a map of Africa illustrating the diverse locations where data was collected to inform the model 2.1. The interaction between temperature and rainfall patterns is crucial in estimating mosquito abundance and distribution. Higher temperatures can increase development rates, while sufficient rainfall provides necessary breeding sites. Regions with favourable combinations of these factors are likely to experience higher mosquito abundance, whereas regions with extreme or insufficient conditions may see reduced mosquito abundance. These interactions, as they occur from the weather data, have been considered in our model.

**Figure 3:**
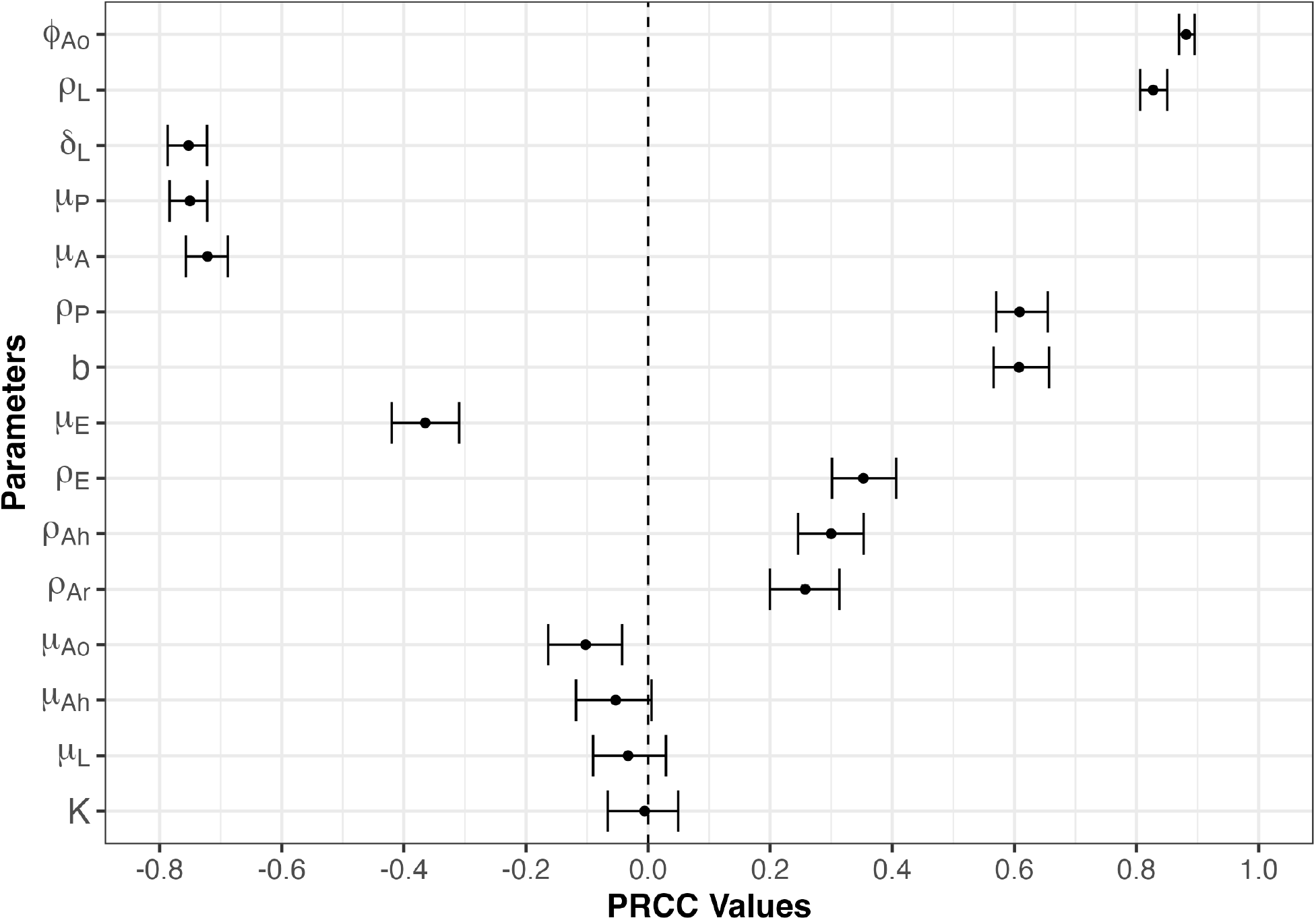
Partial rank correlation coefficients (PRCCs) illustrating the relationship between the total adult mosquito population over time and each model parameter. The PRCCs were derived using 1,000 bootstrap replicates, with input parameters sampled via the Latin hypercube sampling approach. A PRCC value close to 1 indicates a strong positive association, while values near −1 suggest a strong negative association. Parameters with 95% confidence intervals (represented by error bars) overlapping zero are not significantly correlated with the adult mosquito population. Simulations used initial conditions of *E*(0) = 10, *L*(0) = 10, *P* (0) = 10, *A*_*h*_(0) = 10, *A*_*r*_(0) = 10, and *A*_*o*_(0) = 10. These initial conditions differ from those used in the numerical simulations (Section 2.3) to assess the robustness of the sensitivity analysis. No significant differences were observed due to this variation.

The terms in each ODE in system 2.1 correspond to the rate of entry into a stage through the laying or hatching of eggs or maturation and the rate of loss due to maturation and mortality. For further details about the model formulation see Appendix A.1, and for the definition of all variables and parameters see table 1.

**Table 1:** Description of variables and parameters of the model (see Equation (2.1)).

| Variable/<br>Parameter | Description | Unit |
| --- | --- | --- |
| $E(t)$ | Total number of eggs at time t | number |
| $L(t)$ | Total number of larvae at time t | number |
| $P(t)$ | Total number of pupae at time t | number |
| $A_h(t)$ | Total number of host-seeking adult female mosquitoes at time t | number |
| $A_r(t)$ | Total number of adult female mosquitoes at rest for egg development at time t | number |
| $A_o(t)$ | Total number of adult female mosquitoes seeking oviposition site at time t | number |
| $b$ | Total number of eggs laid per oviposition | number |
| $\phi_{Ao}(T)$ | Eggs oviposition rate | day<sup>-1</sup> |
| $\rho_E(T, R)$ | Eggs hatching/development rate | day<sup>-1</sup> |
| $\rho_L(T, R)$ | Larvae development rate into pupae | day<sup>-1</sup> |
| $\rho_P(T, R)$ | Pupae development rate into female adult mosquitoes | day<sup>-1</sup> |
| $\mu_E(T)$ | Egg mortality rate | day<sup>-1</sup> |
| $\mu_L(T)$ | Larvae mortality rate | day<sup>-1</sup> |
| $\mu_P(T)$ | Pupae mortality rate | day<sup>-1</sup> |
| $\mu_A(T)$ | Adult mosquito mortality rate | day<sup>-1</sup> |
| $\mu_{Ao}$ | Adult mortality rate due to oviposition searching site | day<sup>-1</sup> |
| $\mu_{Ah}$ | Adult mortality rate due to host seeking | day<sup>-1</sup> |
| $\delta_L$ | Density-dependent mortality rate of Larvae | day<sup>-1</sup> mosquito<sup>-1</sup> |
| $\rho_{Ah}$ | Rate at which host-seeking adult gets a blood meal and goes to rest | day<sup>-1</sup> |
| $\rho_{Ar}$ | Rate at which adult at rest enter into the oviposition site searching | day<sup>-1</sup> |
| $K$ | Environment carrying capacity of egg-laying female adult mosquitoes | number |

The model 2.1 assumes density-dependent mortality of larvae, accounting for intra- and inter-species larval competition for resources (nutrients) and space (other studies have incorporated density-dependent larval mortality in their models including [12, 26, 32]). A flow diagram of the model 2.1 is shown in figure 1

The model 2.1 is used to understand mosquito stage structure and provide estimates for mosquito abundance due to variability in regional temperature and rainfall. It is an extension of the mosquito population model in Hamdan & Kilicman, (2010) [17], by adding: (i) the effect of temperature and rainfall, (ii) the aquatic stages of the mosquito explicitly, and (iii) density-dependent larval mortality rate. It also extends the non-autonomous mosquito dynamics model in Cailly et al.(2012) [13] and Abdelrazec & Gumel, (2017) [12] by giving a formulation of the temperature- and rainfall-dependent parameters of the model and by incorporating three female adult mosquito stages *A*_*h*_, *A*_*r*_, *A*_*o*_ to provide more accurate population estimates. We use experimental data to parameterize temperature-dependent functions [28, 38, 39]. The functional expressions for rainfall-dependent parameters are extrapolated from the findings in [40], as is also utilized in the investigation outlined in [32].

### 2.2 Parameterization and data

The parameter estimates utilized in the model predominantly rely on data obtained from laboratory studies for *Anopheles* mosquito. To determine the functional relationships in the model, we use nonlinear least squares optimization (NLS) and the method of maximum likelihood estimation (MLE).

For the parameterization of temperature-dependent mortality rates in the juvenile mosquito stages, we drew upon data from two studies conducted by Bayoh & Lindsay [38, 39], which were also used in [28]. These studies investigated mortality at varying constant temperatures using *An. gambiae* mosquitoes, the primary African malaria vector (refer to Appendix A.2). The data indicates a Gaussian dependence of mortality,

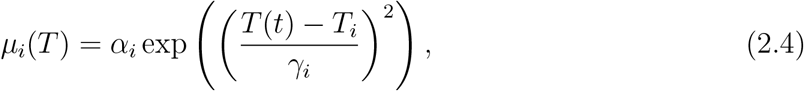

where *i* = (*E, L, P*).

Additionally, the data for temperature-dependent mortality rates in the adult stage originated from a laboratory study to investigate the development and survival of *An. gambiae* at various temperatures [41]. The function that best fits the adult mortality data is,

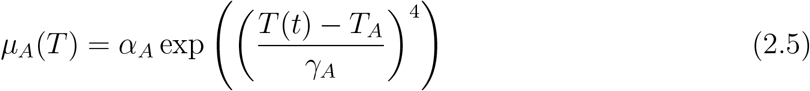

This functional form (equation 2.5) was also used in [28] to model adult mortality. To parameterize the temperature-dependent adult gonotrophic cycle rate, we use data obtained from a research study conducted on *An. pseudopunctipennis* at various constant temperatures [42] (refer to Figure 10a). The data can be represented by

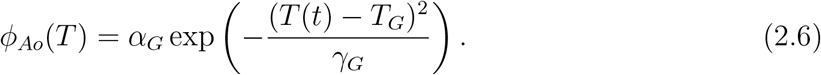

Moreover, to parameterize temperature- and rainfall-dependent developmental rates, we referred to laboratory studies on *Anopheles gambiae* and *Anopheles arabiensis*, as compiled by Depinay et al. [43] and Parham et al. [44]. The data from [44] for the average development times of immature stages *τ*_*i*_ suggest that the functional relation takes the form

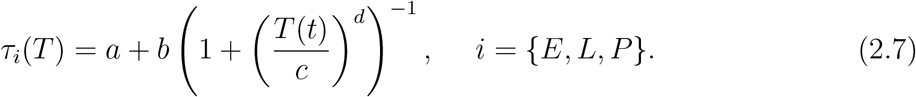

The expression for *τ*_*i*_ follows that which was used in [44] but with slightly different parameter values which were obtained by fitting the function to data to estimate the parameters, *a, b, c*, and *d*.

The maturation rate of eggs into larvae and pupae (into female adult mosquitoes) are defined using the function used in the supplemental material in [44] given by

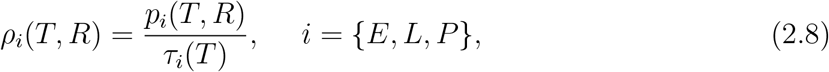

where *τ*_*i*_(*T*) *>* 0 is the average duration of each stage i, given by 2.7 and *p*_*i*_(*T, R*), (where 0 *≥ p*_*i*_(*T, R*) *<* 1) is the daily survival probability of immature mosquitoes in stage i (assumed to be dependent on the mean daily temperature (T^*°*^C) and cumulative daily rainfall (R mm), so that

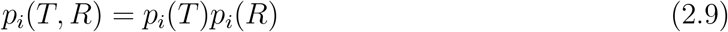

It is worth noting that there may be interactions between temperature and rainfall [28, 45]; however, the nature of the relationship is not clearly studied and therefore we are not integrating this into the model. Hence, we assume independent effects of temperature and rainfall on the daily survival probability of immature mosquitoes (consistent with the assumptions made in [12, 44, 46]). We also assume that development time at each stage is dependent only on temperature if there is adequate rainfall to sustain development [44]. Following [44, 46], the daily survival probability of immature mosquitoes in stage i due to temperature is given by

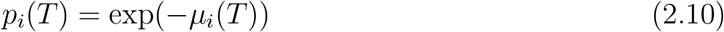

Also, following the study in the supplementary material of [44] and the work done in [46], the rainfall-dependent daily survival probability of immature mosquitoes, *p*_*i*_(*R*) is given by

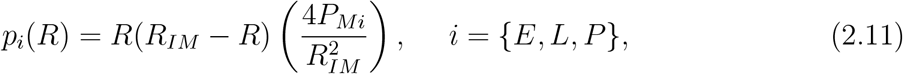

where *P*_*Mi*_ is the peak daily survival probability of immature mosquito in stage i and *R*_*IM*_ *> R*(*t*) *>* 0, for all t, is the rainfall threshold for a particular region beyond which wash out effects from excessive rainfall cause no immature mosquitoes to survive.

The aforementioned parameterized functions given by equations 2.4, 2.5 and 2.6 are plotted in figure 10 in the Appendix. For the development rate functions (equation 2.8), we use mean monthly temperature and rainfall data from Ain Mahbel, Algeria to produce the figures (see the appendix 4 for details describing the data). Numerical simulations of the model are carried out using temperature and rainfall profiles of four cities in East, West, North, and South Africa (see figure 2). These regions exhibit diverse seasonality patterns, ranging from year-round mosquito presence in Kumasi, Ghana, and Nairobi, Kenya, to a markedly seasonal mosquito abundance in Ain Mahbel, Algeria, and Cape Town, South Africa [47]. The primary mosquito species utilized for gathering data to estimate parameters are *Anopheles* mosquitoes; however, this model can be re-parameterized and used to study various other mosquito species.

### 2.3 Numerical Analysis

The numerical solutions of the model (2.1) are obtained using the deSolve package in R. This investigation aims to explore the impact of local environmental conditions, specifically temperature and rainfall, on mosquito population dynamics and stage structure. The parameterized functions detailed in Section 2.2 and the constant parameters outlined in Table 1 and 2 are used for the analysis of the model. We assume initial conditions (*E*(0) = 100.0, *L*(0) = 100.0, *P* (0) = 100.0, *A*_*h*_(0) = 30.0, *A*_*r*_(0) = 30.0, *A*_*o*_ = 40.0) for the numerical simulations.

**Table 2:** Parameter values and ranges used in the mosquito population model (2.1). The ranges were selected to reflect average monthly temperatures of 15–34°C and rainfall levels of 10–300 mm.

| Parameter | Range | Baseline Value | References |
| --- | --- | --- | --- |
| $b$ | [150, 300] | 200 | [12, 48] |
| $\delta_L$ | [0.02, 0.06] | 0.03 | [27] |
| $\rho_{Ah}$ | [0.6, 2.3] | 2 | [13, 49] |
| $\rho_{Ar}$ | [0.5, 1.5] | 0.5 | [13, 49] |
| $K$ | [500, $3 \times 10^8$] | $10^8$ | [23, 27, 50] |
| $\mu_{Ao}$ | [0.20, 0.45] | 0.25 | [26] |
| $\mu_{Ah}$ | [0.12, 0.23] | 0.18 | [26] |
| $T\phi_{Ao}$ | [0.03, 0.45] | 0.24 | [28] |
| $T\mu_E$ | [0.06, 0.24] | 0.15 | [28, 44] |
| $T\mu_L$ | [0.025, 0.10] | 0.06 | [28, 44] |
| $T\mu_P$ | [0.05, 0.24] | 0.15 | [28, 44] |
| $T\mu_A$ | [0.075, 0.13] | 0.10 | [27, 44, 51] |
| $T\rho_E$ | [0.17, 0.5] | 0.335 | [12, 27, 44, 46] |
| $T\rho_L$ | [0.05, 0.1] | 0.09 | [12, 27, 44, 46] |
| $T\rho_P$ | [0.14, 0.33] | 0.235 | [12, 27, 44, 46] |

### 2.4 Sensitivity Analysis

To assess the influence of model parameters on the total adult mosquito population, we conducted a global sensitivity analysis using the Latin hypercube sampling (LHS) method combined with partial rank correlation coefficients (PRCC) [23, 26, 52, 53, 54]. The adult mosquito stage was selected as the primary output variable due to its epidemiological importance as the vector stage for mosquito-borne diseases, with many intervention strategies aimed at reducing this population [55, 56, 57]. We assumed a uniform probability distribution for each parameter within the ranges specified in Table 2, covering average monthly temperatures and cumulative rainfall between 15–34°C and 10–300 mm, respectively. This uniform distribution allows each parameter value an equal probability within its range, enabling comprehensive exploration of the parameter space [54, 58].

The mosquito model (2.1) was solved numerically for a latin hypercube sampled set of parameter values (n = 1000), and PRCC values were calculated between the total adult mosquito population and each parameter, using 1000 bootstrap replicates to estimate confidence intervals. PRCC values range from −1 to 1, where values near 1 indicate a strong positive correlation with the output variable, values near −1 indicate a strong negative correlation, and values near 0 imply limited influence on the output variable. All analyses were conducted using the “lhs,” “sensitivity,” and “boot” packages in R [59, 60, 61].

## 3 Results

In our analysis of the four considered regions in Africa, it became evident that neglecting seasonality leads to significant overestimation or underestimation of mosquito abundance. Figure 4 illustrates the comparison between constant and seasonal environmental conditions. For Ain Mahbel, Algeria, disregarding seasonality results in an underestimated mosquito population. However, when seasonality is taken into account, we observe a substantial mosquito population, particularly during June to October. Conversely, in Ghana, neglecting seasonality leads to an overestimation of the total mosquito abundance for the year compared to when seasonality is considered. Specifically, in Algeria, the estimated annual mosquito abundance is approximately 11,434 when seasonality is ignored. This number is about ten times less than the population estimated when seasonality is considered.

**Figure 4:**
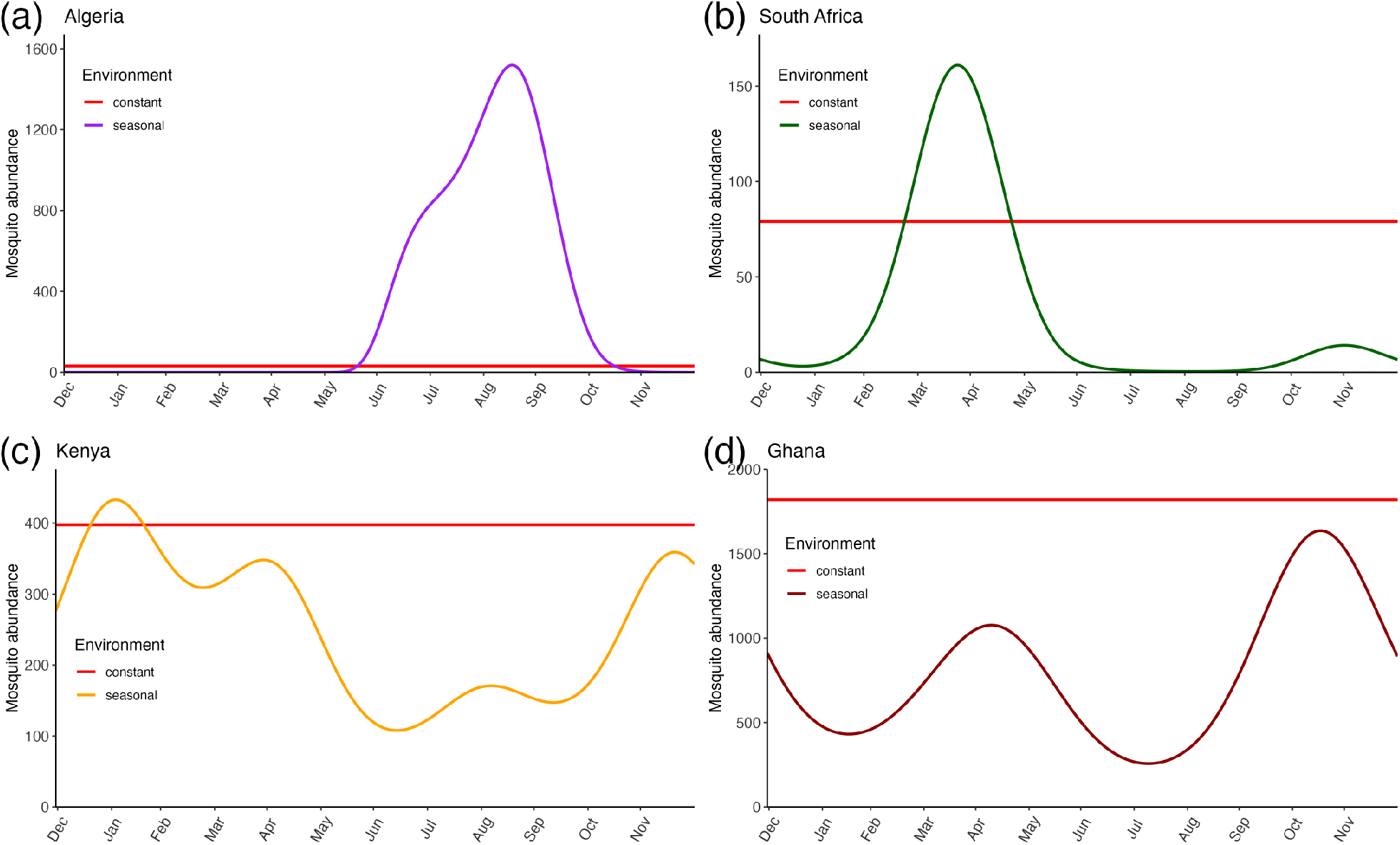
Numerical simulation of the model 2.1 illustrating the abundance of female adult mosquitoes in four African cities, comparing the population dynamics under constant and seasonal environmental conditions. Constant environmental condition refers to the averaging of temperature and rainfall conditions, where monthly temperature and rainfall values were averaged across the year and kept constant, and represents the parametrization approach that is used for models that do not consider seasonality. Seasonal environmental condition refers to the natural variations in temperature and rainfall throughout the year, reflecting the actual seasonal changes experienced in each region. These conditions were applied uniformly across all regions studied to ensure comparability of results. (a) Ain Mahbel, Algeria (*R*_*IM*_ = 60 mm), (b) Kumasi, Ghana (*R*_*IM*_ = 550 mm), (c) Nairobi, Kenya (*R*_*IM*_ = 200 mm), and (d) Cape Town, South Africa (*R*_*IM*_ = 250 mm). These figures present results from the 5th-year simulation to mitigate initial condition effects.

We find that in different regions mosquitoes have different population dynamics (Figure 5). The region with warmer temperatures and higher rainfall (Kumasi, Ghana) across the year seems to have higher mosquito abundance (Figure 5b). Conversely, the region with cold temperatures and moderate rainfall (Cape Town, South Africa) has a much smaller mosquito abundance and peaks once for the entire year with mosquito numbers averaging around 11,426 which is about 25 times less than the estimated value in Ghana. The cold weather give rise to declines in the number of larvae that emerge as adults but these abundances rebound when environmental conditions become favourable as seen in Figure 5c-f. The smaller population size during the months of November-May in Algeria and June-January in South Africa is most likely due to the cold temperatures (Figure 5c-f) resulting in longer juvenile development times leading to a small number of larvae emerging as adults. Figure (5c-f) shows that temperature and rainfall may greatly impact mosquito stage structure.

**Figure 5:**
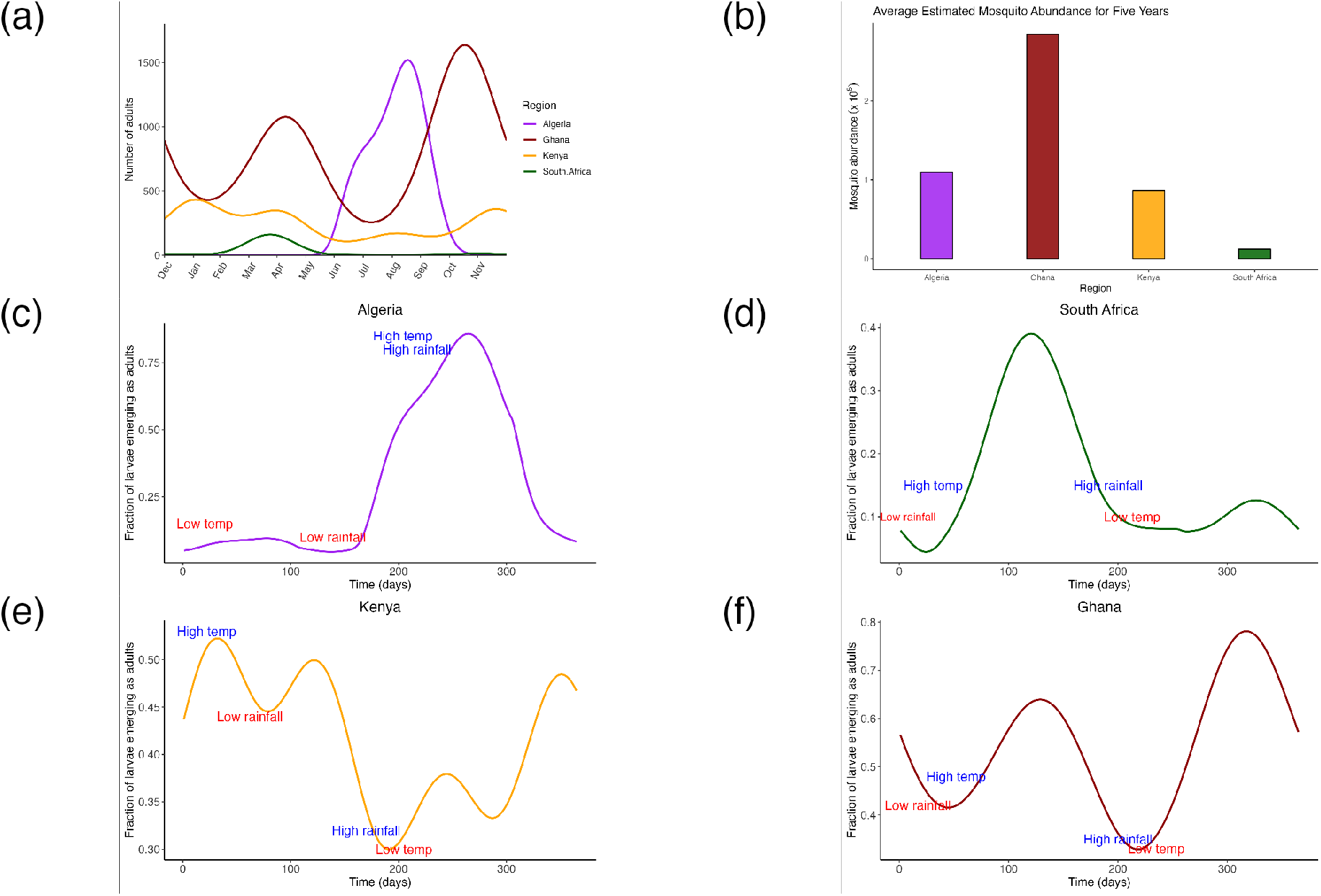
Numerical solutions of the stage-structured model. All figures present results from the 5th year of the simulation as by this time, dynamics with a year are not affected by the initial conditions. (a) Monthly dynamics of adult female mosquito populations compared across four African cities. (b) Yearly total abundance estimates for each region. (c) Daily fraction of larvae emerging as adult mosquitoes in Algeria. (d) Daily fraction of larvae emerging as adult mosquitoes in South Africa. (e) Daily fraction of larvae emerging as adult mosquitoes in Kenya. (f) Daily fraction of larvae emerging as adult mosquitoes in Ghana.

Figure 6 shows that mosquito abundance may vary across different months between regions. The model describing the region with warmer temperatures and higher rainfall (Kumasi, Ghana) estimates two peaks in mosquito abundance (that are in May and November) when temperature and rainfall values are in the range [26-28]^*°*^C and [104-230]mm respectively. The region with colder temperatures, Algeria, exhibits one peak during September when temperature and rainfall values are 23.5^*°*^C and 62mm. In Kenya, characterized by relatively mild and stable environmental conditions with less pronounced seasonality (see Figure 2), we observe a substantial presence of mosquitoes throughout the year, with a significant fraction of annual mosquito abundance observed each month. Each of the regions peaks at different months and different levels. This implies that the timing of mosquito control implementation may differ for different regions.

**Figure 6:**
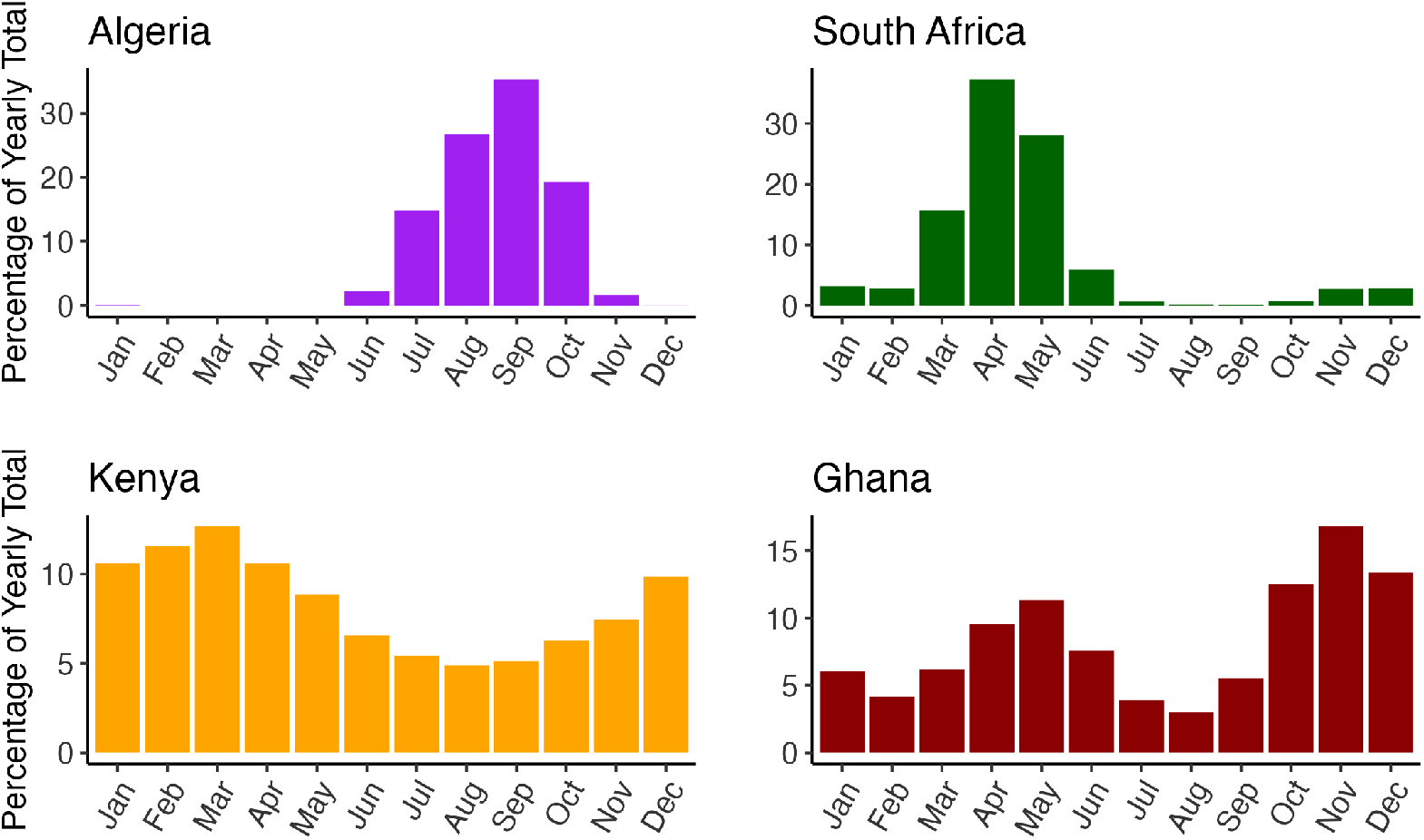
Model simulations illustrating the monthly distribution of adult female mosquitoes as a percentage of the yearly total across regions. Results are averaged over a five-year simulation period.

To assess the optimal timing and duration of mosquito control interventions throughout the year, we calculated the cumulative fraction of mosquitoes for each region (see Figure 7a). Our findings reveal that in regions with colder temperatures, such as Algeria and South Africa, approximately 50% of the annual mosquito population can be controlled by focusing interventions implemented during just two months. For instance, in Algeria, targeting control measures during August and September could account for half of the mosquito population for the entire year. Conversely, in warmer regions, such as Kenya, vector control efforts may need to be intensified over a longer period, spanning approximately 4 to 5 months, to achieve a similar level of control. Our analysis suggests that in Kenya, implementing controls from December to May could cover 50% of the annual mosquito abundance. Furthermore, our results indicate variations in the optimal timing of control strategies between regions, as illustrated in Figure 7b. Specifically, regions like Ghana and Kenya may require continuous vector control efforts throughout the year, whereas in Algeria and South Africa, vector controls may be concentrated within a few months.

**Figure 7:**
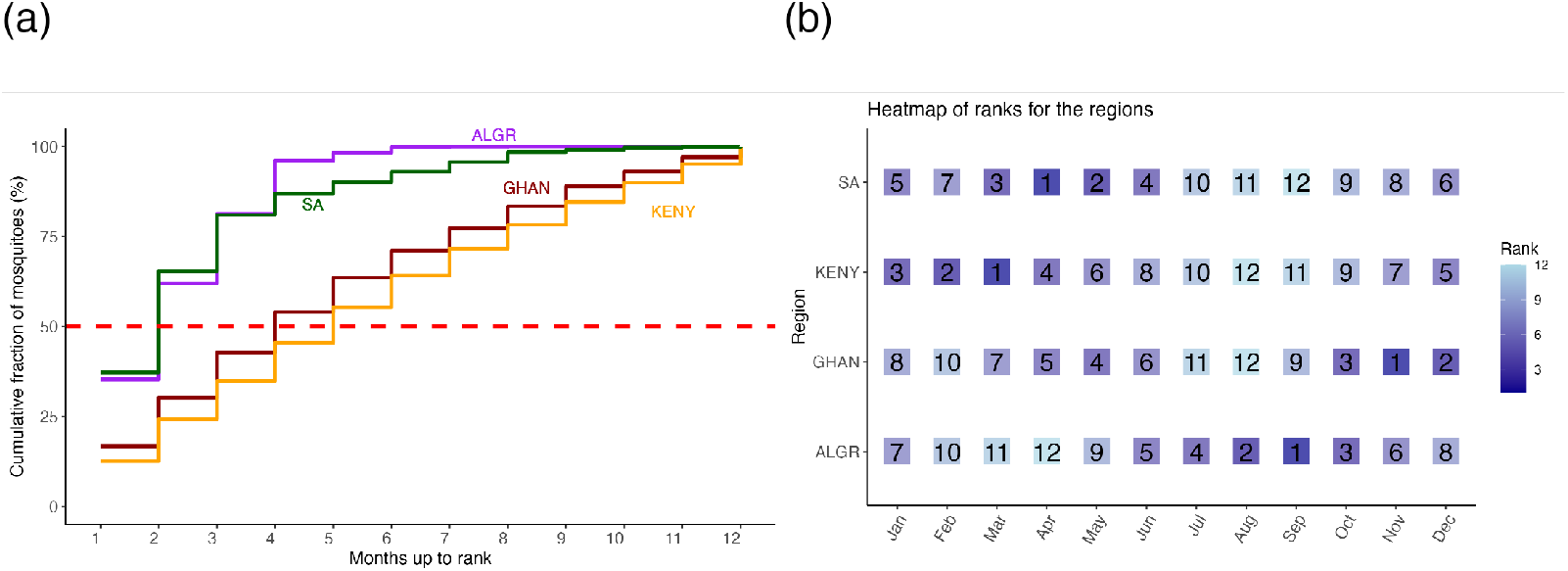
Examining the optimal timing for mosquito control strategies, we calculated the cumulative fraction of mosquitoes for each region. This analysis highlights specific months requiring concentrated control efforts to effectively reduce mosquito populations. Notably, in Algeria and South Africa, over half of the mosquito population is concentrated within two months. Focusing control measures during these months could significantly diminish mosquito populations and consequently mitigate the spread of diseases.

Figure 8 is a comparison between mosquito abundance and human deaths attributed to malaria across the various regions from 2015 to 2019. Mosquito abundance data, generated from the model 2.1, is compared with malaria-related mortality data obtained from the Institute for Health Metrics and Evaluation (IHME), Global Burden of Disease (2019) [62] as a check on whether the model’s predictions are broadly consistent with observations. Notably, regions with higher mosquito abundance, such as Ghana, consistently exhibit elevated levels of malaria-related deaths during the study period (see Figure 8). Conversely, regions with lower mosquito populations correspondingly report fewer malaria deaths. However, Algeria presents a distinct pattern; despite estimating a significant mosquito population, no malaria-related deaths are reported during this period. This anomaly can be attributed to Algeria’s official recognition by the World Health Organization (WHO) as malaria-free, making it the second country in the WHO African region to achieve this status. Algeria recorded its last indigenous malaria cases in 2013 [63] which is before the period considered in this study.

**Figure 8:**
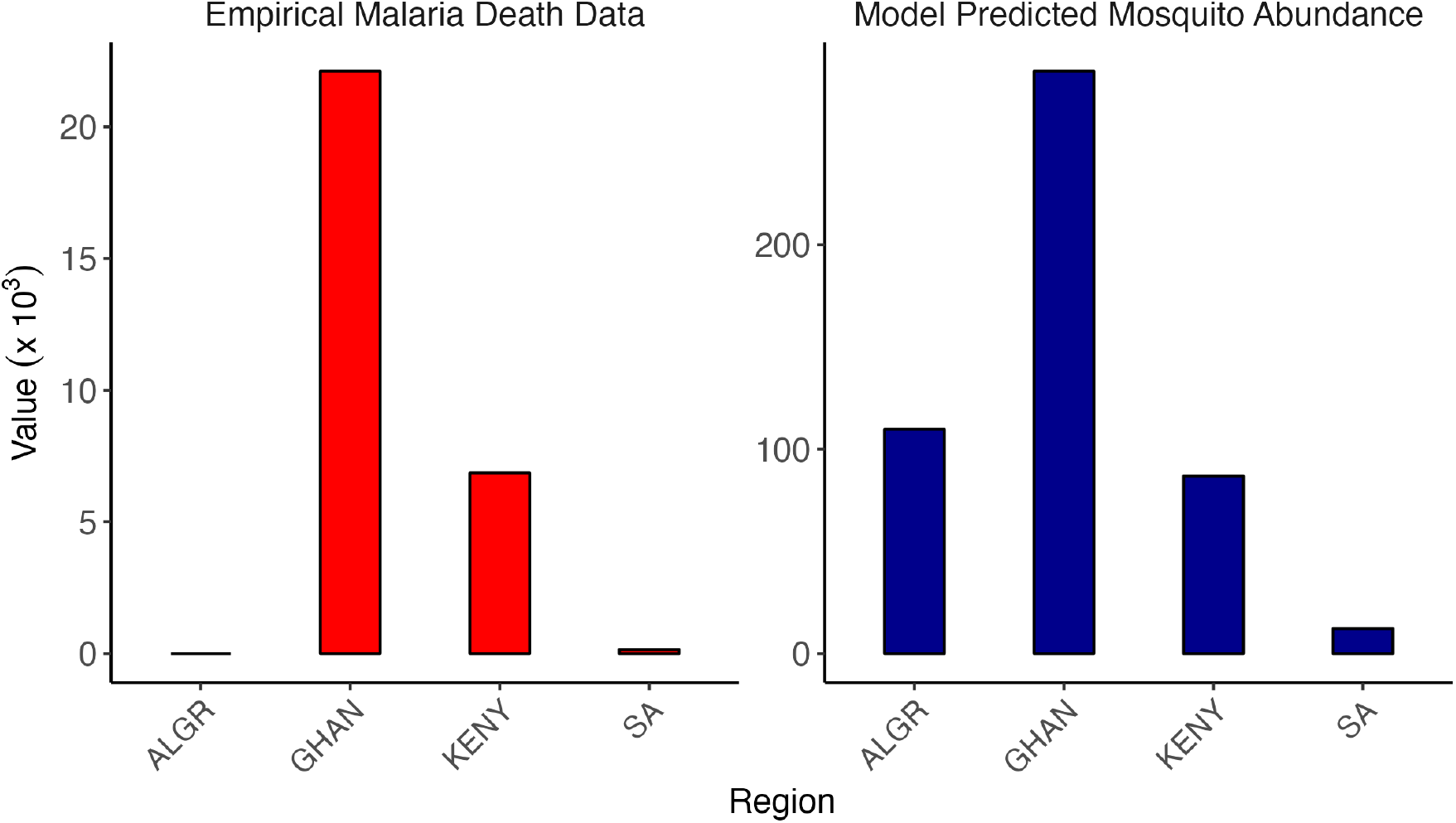
Comparison of estimated mosquito abundance and human malaria-related deaths across four regions from 2015 to 2019. Malaria motality data were obtained from IHME, Global Burden of Disease (2019).

To assess performance, the model 2.1 is compared with *Anopheles* species abundance data collected from different sites within the regions under consideration. It is important to note that the environments for the various regions where mosquitoes were collected are not free from vector control interventions, which may affect the number of mosquitoes collected. Further details on mosquito collection and experimental procedures for each region can be found in [64, 65] for Ghana, [66] for Kenya, and [67] for South Africa. In Figure 9, we show the comparison of estimated mosquito abundance to actual mosquito counts for the different regions. We find that in one site in South Africa, there is a close correlation between the model estimates and the actual mosquito count data (Figure 9a) (the model produces a similar curve with the mosquito count data) except that the peak times differ. For Ghana, the model agrees with the data at some points and it deviates at some other points as shown in Figure 9c. For Kenya, the data for the different sites do not agree with the model’s estimates. This may be due to data collection issues, such as low sample size, vector control implemented that reduces the number of vectors available to be trapped and counted, or it may indicate that model assumptions are violated and that our modelling framework needs to be further refined.

**Figure 9:**
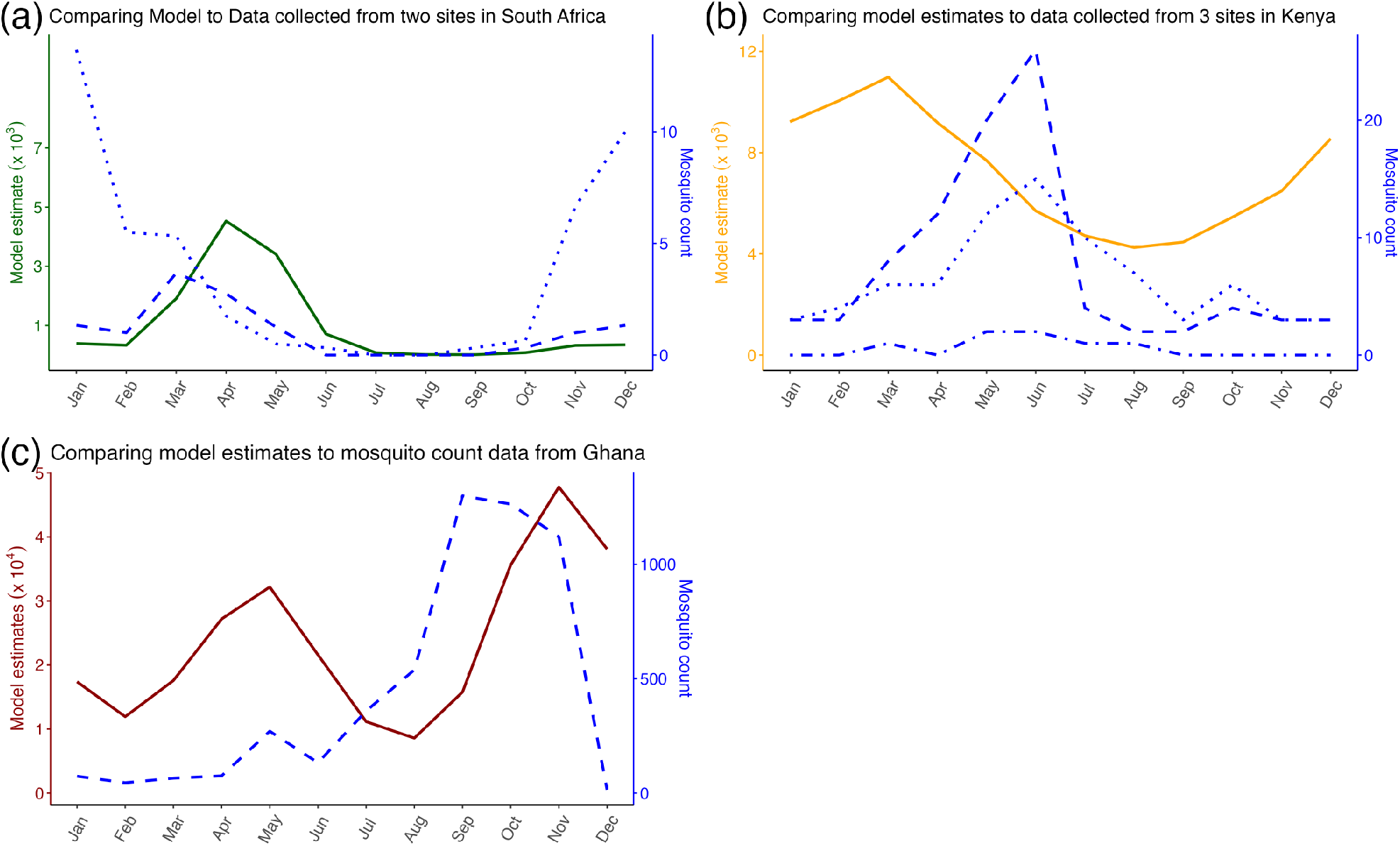
Comparing average *Anopheles* mosquito count data with simulated monthly mosquito abundance for (a) South Africa, (b) Kenya, and (c) Ghana.

**Figure 10:**
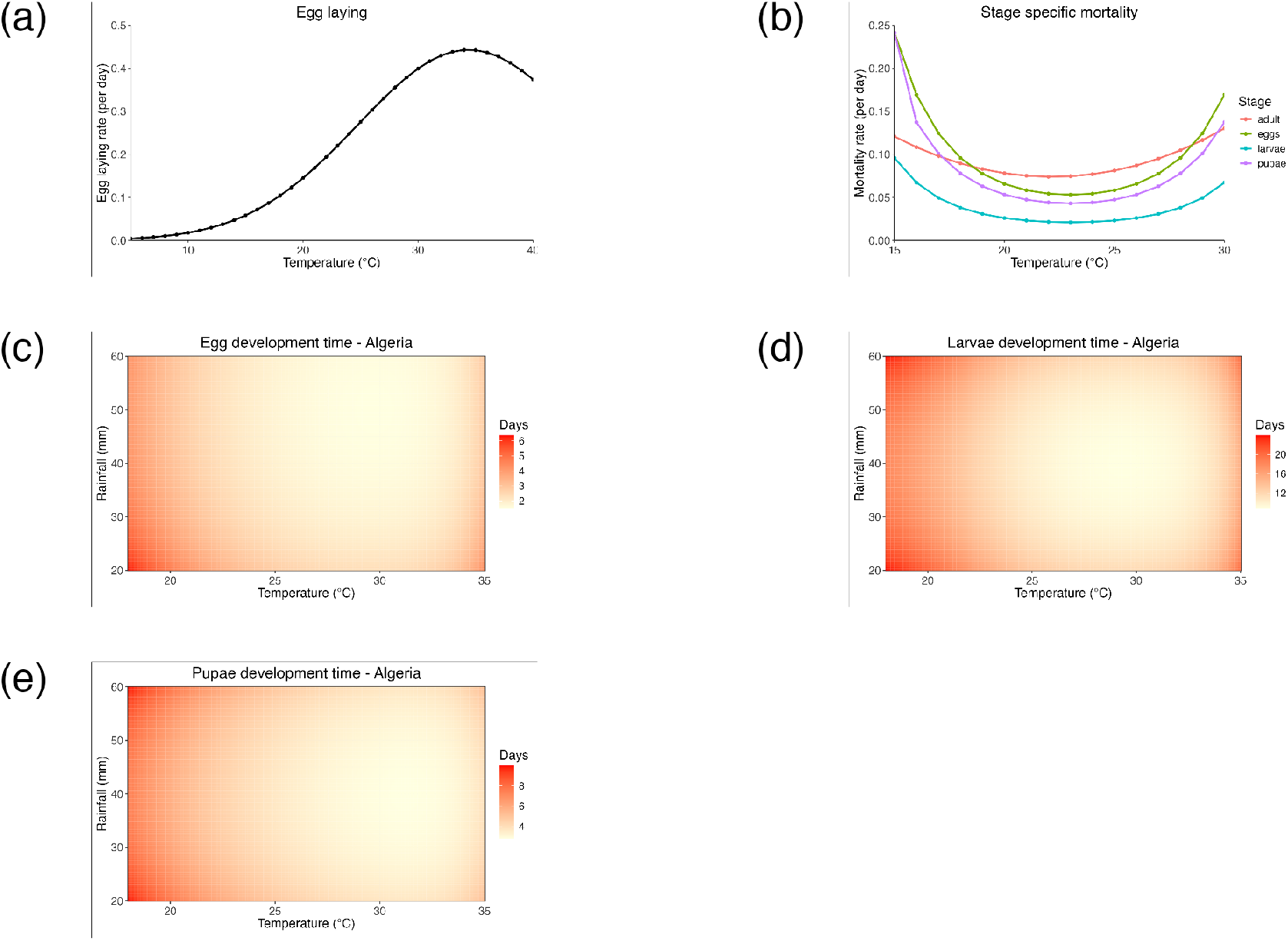
Functional relationships used to describe the relationship between life history parameters and weather variables including temperature and rainfall.

In our sensitivity analysis, we found that the egg oviposition rate of adult mosquitoes (*ϕ*_*Ao*_) and the larvae development rate (*ρ*_*L*_) exhibited strong positive correlations with the adult mosquito population (PRCC *>* 0.7; see Figure 3). Additionally, the pupae development rate into adult mosquitoes (*ρ*_*P*_) (PRCC = 0.608), the number of eggs laid per oviposition (PRCC = 0.607), the egg hatching rate (*ρ*_*E*_) (PRCC = 0.35), the rate at which host-seeking adults obtain blood meals (*ρ*_*Ah*_) (PRCC = 0.29), and the rate at which resting adults transition to oviposition site searching (*ρ*_*Ar*_) (PRCC = 0.25) demonstrated moderate positive correlations. Conversely, strong negative correlations were observed between the adult mosquito population and the density-dependent mortality rate of larvae (*δ*_*L*_) (PRCC = −0.75), pupae mortality rate (*µ*_*P*_) (PRCC = −0.75), and adult mosquito mortality rate (*µ*_*A*_) (PRCC = −0.72). The mortality rate of adults due to oviposition site searching (*µ*_*Ao*_) showed a weak negative correlation (PRCC = −0.10). Other parameters, including the adult mortality rate due to host seeking, larvae mortality rate, and the environmental carrying capacity for egg-laying female adults (*K*), had PRCC values with 95% confidence intervals overlapping zero, indicating they do not significantly influence the adult mosquito population. These findings align with the results of the sensitivity analysis presented in [12].

## 4 Discussion

Our findings underscore the importance of considering regional seasonality in weather for accurate modelling of mosquito population dynamics and stage structure. Understanding these dynamics is crucial for effective vector-borne disease outbreak prediction and vector management strategies in different regions. The analysis reveals that incorporating seasonality in environmental conditions is essential, as neglecting it may lead to biased predictions, as shown in Figure 4. Under constant temperature and rainfall, the mosquito population reaches a stable equilibrium where birth and death rates balance perfectly, resulting in no temporal fluctuations. This behavior stems from the deterministic nature of the model, which assumes constant resources and fixed biological rates under constant conditions. While this simplifies the dynamics, real-world systems often exhibit stochastic fluctuations due to random ecological and biological processes. The inclusion of constant environmental conditions in our study serves as a baseline to highlight the critical role of seasonality in driving mosquito abundance. Therefore, integrating seasonality into models is imperative for producing reliable predictions and supporting informed decision-making in vector control efforts.

The dynamics of mosquitoes in regions with colder temperatures, such as Algeria and South Africa, exhibit similarities, characterized by large peaks during favourable environmental conditions with reduced populations during other periods (Figure 5a). Conversely, in relatively warmer regions like Ghana, mosquitoes persist throughout the year, with consistently high population estimates. Kenya experiences a moderate range of temperatures and varied rainfall, leading to mosquito persistence at lower levels compared to Ghana. Regions with colder temperatures, such as Algeria and South Africa, experience population declines during colder months, followed by rebounds and population increases during warmer periods (Figure 5a, c, d). These seasonal patterns significantly influence the dynamics of mosquito populations (either increasing or decreasing mosquito abundance depending on how favourable environmental conditions are) and, consequently, the transmission cycles of mosquito-borne diseases.

Our analysis reveals distinct patterns in mosquito abundance across different months and regions, as depicted in Figure 6. Regions experiencing warmer temperatures and higher rainfall, like Kumasi, Ghana, exhibit two peaks in mosquito abundance in May and November, coinciding with optimal environmental conditions for mosquito development. During these months, reduced development times lead to accelerated juvenile maturation to the adult stage. Conversely, colder regions such as Algeria show a single peak in September. Interestingly, regions like Kenya, with milder and less seasonal environmental conditions, maintain a consistent presence of mosquito vectors throughout the year. These findings suggest the importance of tailoring mosquito control efforts to specific environmental conditions. In colder regions like Algeria and South Africa, focusing vector management efforts during peak periods could significantly reduce mosquito populations and control disease spread. However, the effectiveness of control measures may vary due to regional environmental variations. Therefore, adaptive and context-specific vector control strategies are essential to reduce mosquito populations and mitigate mosquito-borne diseases effectively.

Our analysis reveals varying durations of control implementation across regions, reflecting differences in local environmental conditions. Regions with warmer temperatures and higher rainfall exhibit higher mosquito populations year-round, necessitating continuous vector control efforts. Conversely, regions with colder temperatures experience minimal mosquito populations during certain periods, requiring control measures only during select months. This insight underscores the importance of identifying critical periods for mosquito control to effectively reduce populations and mitigate disease spread.

The analysis of malaria mortality data underscores a significant agreement between mosquito abundance and malaria deaths during the period from 2015 to 2019 (see Figure 8). Ghana, estimating the highest mosquito populations throughout the year among the regions studied, also reports the highest malaria deaths. Conversely, Algeria, despite estimating a substantial mosquito population, reports zero malaria deaths during the same period. This anomaly is attributed to Algeria’s achievement of malaria-free status, certified by the World Health Organization (WHO) during this time [63]. Algeria’s success in combating the disease can be attributed to various factors, including a well-trained health workforce, universal health-care providing malaria diagnosis and treatment, prompt responses to disease outbreaks, and vector control efforts. Algeria’s example provides valuable lessons for other regions in Africa, demonstrating that malaria elimination is attainable through concerted efforts, strategic investments, and evidence-based approaches.

Upon comparing the model-derived estimates with actual mosquito collection data (Figure 9) to assess model performance, a notable lack of agreement between the two sets of data and the model prediction becomes apparent. This discrepancy may stem from biases and under-sampling of mosquitoes at collection sites, or that data to test the model are insufficient. However, Figure 9 may also indicate that model assumptions are too simplified to quantitatively predict mosquito abundance, and for this reason many of the model results have been stated as qualitative characteristics such as the number of peaks in adult abundance and the within-year variability.

The sensitivity analysis as discussed in section 2.4 highlights the critical parameters influencing the adult mosquito population and underscores the importance of targeting these parameters for effective control strategies. As shown in Figure 3, the most influential parameters include the oviposition rate of adult mosquitoes (*ϕ*_*Ao*_), larvae development rate (*ρ*_*L*_), pupae development rate into adult mosquitoes (*ρ*_*P*_), number of eggs laid per oviposition (*b*), density-dependent mortality rate of larvae (*δ*_*L*_), pupae mortality rate (*µ*_*P*_), and adult mosquito mortality rate (*µ*_*A*_). Parameters such as *ϕ*_*Ao*_, *ρ*_*L*_, *ρ*_*P*_, and *b* are positively correlated with the adult mosquito population, while *δ*_*L*_, *µ*_*P*_, and *µ*_*A*_ exhibit negative correlations. This suggests that control strategies designed to reduce *ϕ*_*Ao*_, *ρ*_*L*_, *ρ*_*P*_, and *b*, while simultaneously increasing *δ*_*L*_, *µ*_*P*_, and *µ*_*A*_, could significantly diminish mosquito abundance in a region. For instance, larvicides and adulticides can decrease resources available for larvae development, thereby intensifying competition among immature mosquitoes, which leads to higher density-dependent mortality (*δ*_*L*_) and increased pupae mortality (*µ*_*P*_). These interventions also reduce the development rate of pupae into adults (*ρ*_*P*_). Additionally, the use of personal protective measures such as insecticide-treated bed nets, insect repellents, and indoor residual spraying can effectively lower the oviposition rate (*ϕ*_*Ao*_) and reduce the number of eggs laid (*b*), ultimately suppressing mosquito population growth.

It’s important to highlight that our model formulation does not explicitly account for adult males. Instead, it assumes that their abundance is equal to that of adult females. This simplification is rooted in the epidemiological significance of female mosquitoes. Unlike their male counterparts, female mosquitoes serve as vectors responsible for transmitting pathogens because they require a blood meal for egg production. While males play a role in reproduction, they do not contribute to disease transmission as they do not feed on blood. Therefore, for epidemiological relevance, assuming equal abundance between adult males and females is justified. This simplification helps streamline the model’s complexity without compromising its accuracy in investigating mosquito dynamics.

While our model provides valuable insights into mosquito population dynamics and their response to environmental conditions, several limitations should be noted. Firstly, the model simplifies certain biological processes by not explicitly considering adult male mosquitoes in the formulation. While we assume equal abundance between adult males and females, this simplification may overlook potential differences in behaviour and ecology between the sexes that could influence population dynamics. Additionally, our model primarily focuses on the *Anopheles* species, a significant malaria vector, potentially neglecting the dynamics of other mosquito species present in the study regions, such as the *Aedes* mosquito in South Africa, and their interactions with *Anopheles*. It is important to note that the potential presence of other mosquito species leading to intra-specific competitions may impact local dynamics (as discussed in [11]). Furthermore, the model assumes homogeneity in environmental conditions within each region, disregarding potential micro-climatic variations that could impact mosquito population dynamics.

Another limitation of our model is the assumption that mosquito mortality rates are primarily temperature-dependent, while the direct effects of rainfall on mortality are not explicitly modelled due to a lack of empirical evidence. Although we account for the impact of rainfall indirectly through its effect on development rates, further research is needed to better understand and quantify the relationship between rainfall and mosquito mortality. Moreover, while temperature and rainfall data are incorporated into the model, other important factors like habitat suitability, land use changes, and human interventions (e.g., vector control measures) are not explicitly considered, despite their potential impact on mosquito populations. The accuracy of our model estimates is also dependent on the quality of input data, including parameter estimates and environmental variables, which may vary in availability and reliability across regions. Additionally, the use of monthly averaged temperature and rainfall data may not adequately capture micro-fluctuations in environmental conditions, as daily data were not readily available for most study regions. Furthermore, the model fit to temperature and rainfall data was not good in some regions (see figure 2 and table 3), potentially affecting the accuracy of the model estimates. Therefore, while our model offers valuable insights, further refinements and validations are necessary to improve its robustness and applicability.

**Table 3:** *R*^2^ values for function fittings to temperature and rainfall data across four African regions.

| Region | Temperature (T(t)) | Rainfall (R(t)) |
| --- | --- | --- |
| Ghana | 0.97 | 0.62 |
| Algeria | 0.99 | 0.96 |
| Kenya | 0.90 | 0.64 |
| South Africa | 0.99 | 0.88 |

Our study investigates the influence of local temperature and rainfall conditions on mosquito population dynamics across diverse regions.Our model is specifically designed to capture the intricate behaviour of the *Anopheles* species, a key vector responsible for malaria transmission and prevalent in the regions under consideration. Through our model analysis, we obtain insights into the intricate interplay between environmental factors and mosquito population dynamics, thereby enriching our understanding of optimal timing for vector control implementation and vector-borne disease transmission dynamics.

## 5 Conclusion

This study formulates a temperature- and rainfall-dependent stage-structured model, shedding light on the influence of regional weather variability on mosquito population dynamics and stage structure across Ain Mahbel, Algeria, Cape Town, South Africa, Nairobi, Kenya, and Kumasi, Ghana. The observed variations in mosquito population dynamics among these regions emphasize the need to account for local weather conditions in vector control planning.

The identification of peak mosquito months provides invaluable insights for the formulation of targeted intervention strategies. Moreover, the relationship between mosquito dynamics and malaria-related deaths underscores the potential implications for disease outbreak risk and duration. The sensitivity analysis further highlights the critical role of parameters such as oviposition rate, development rates, and mortality rates in shaping mosquito population dynamics, offering insights for designing effective control measures. These findings significantly contribute to our broader comprehension of vector-borne disease dynamics and provide a foundation for the development of region-specific vector control strategies aimed at mitigating outbreak risks.

Future research avenues may extend the scope of this study to include other regions where malaria is endemic. Additionally, there is a need to explore the integration of additional environmental factors such as humidity to obtain a more comprehensive understanding of the impact of environmental conditions on mosquito population dynamics and the risk of disease transmission. Furthermore, future studies could delve deeper into sensitivity analyses to understand how changes in key parameters affect model predictions, enhancing the model’s credibility and reliability while refining intervention strategies.

## Acknowledgements

Amy Hurford acknowledges financial support from the Natural Sciences and Engineering Research Council of Canada (NSERC) through Discovery Grant RGPIN-2023-05905. The authors gratefully acknowledge the valuable feedback provided by members of the Hurford lab.

## Statements and Declarations

Amy Hurford acknowledges financial support from the Natural Sciences and Engineering Research Council of Canada (NSERC) through Discovery Grant RGPIN RGPIN-2023-05905.

## A Model description and parameterization

### A.1 Model description

Here we derive the mathematical model describing mosquito dynamics (see equation 2.1). This model 2.1 is very similar to the models formulated in Abdelrazec & Gumel, (2017) [1] and also the model in Caily et. al., (2012) [2]. The major difference between our model and that of Abdelrazec & Gumel, (2017) [1] is that our model has the adult population subdivided into three stages (*A*_*h*_(*t*), *A*_*r*_(*t*), *A*_*o*_(*t*)). Also comparing our model to Caily et. al., (2012), our model has the parameters depending on time.

Our mathematical model for mosquito population dynamics incorporates a temperature- and rainfall-dependent development rates and temperature-dependent mortality rates. We also incorporates density-dependent mortality in the laval stage. We note that temperature and rainfall are time-dependent.

Let *E*(*t*), *L*(*t*), *P* (*t*) and *A*(*t*) be the distinct life stages of the mosquito. Assuming that the adult females mate with the male counterparts just upon emerging from the pupa, we describe a stage-structured mathematical model as;

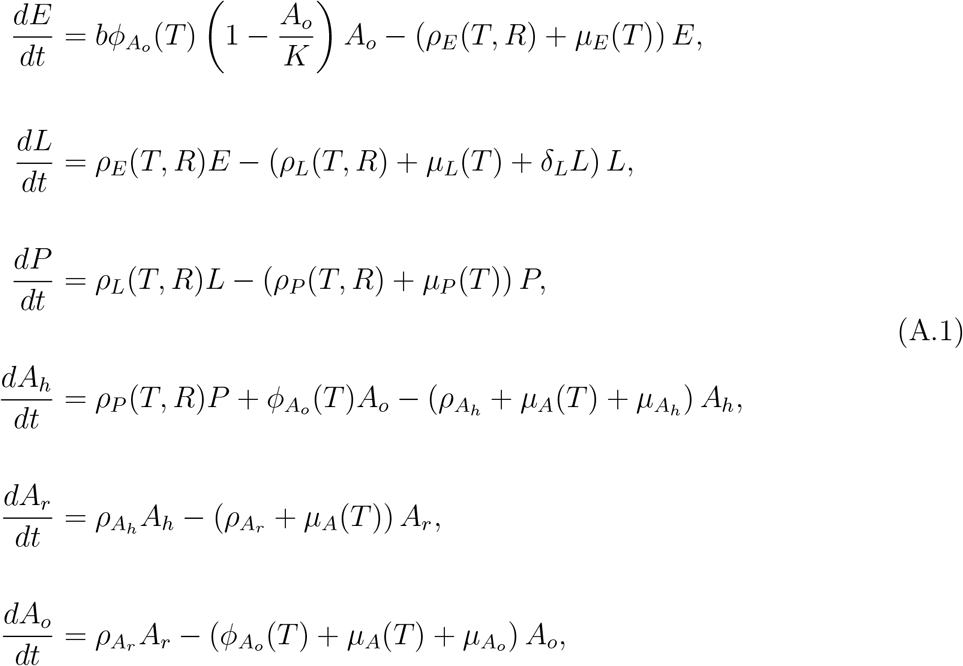

The term 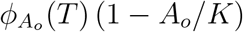 represents the density-dependent eggs oviposition rate (where *K > A*_*o*_) for all *t ≥* 0 is the environmental carrying capacity of female adult mosquitoes and 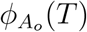 is the temperature-dependent egg deposition rate. Eggs hatch into larvae stage at a temperature- and rainfall-dependent rate, *ρ*_*E*_(*T, R*). Larvae mature into pupae at a temperature- and rainfall-dependent rate, *ρ*_*L*_(*T, R*). Pupae mature into the female adult mosquitoes at a temperature- and rainfall-dependent rate, *ρ*_*P*_ (*T, R*). It should be stated that the development rates for the juvenile stages are dependent on temperature and rainfall because, while temperature values greatly impact developmental time of the juvenile mosquitoes [3, 4, 5, 6, 7, 8], rainfall is required for availability of breeding sites and habitats [8, 9, 10]. However, extreme environmental conditions such as too hot or too cold are not favourable for the survival and maturation of the larvae [3, 4, 5, 6, 8, 11] and also excessive rainfall washes out the larval breeding sites, such as small stagnant water on yards or lawn [1]. These newly emerged female adult mosquitoes mate with the males and quest for a blood meal. Upon successful feeding on a blood meal, they go to rest to allow for the eggs to develop at a rate 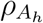. The rested female with developed eggs seeks for oviposition site at a rate 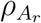. Ones the eggs are laid, it goes through the cycle again and the mosquito which laid the eggs also go into the adult seeking host stage to continue the cycle until it dies and leaves the cycle.

### A.2 Data and parameter estimation

Description of all parameters are provided in table 1 in the main manuscript. Regional variations exist in temperature and rainfall, yet the functional relationships describing the model parameters with respect to temperature (T) and rainfall (R) remain consistent across diverse regions. The estimation of life history parameters as a function of temperature and/or rainfall involves curve fitting to laboratory experiment data utilizing functional forms based on previous studies [8, 10, 12, 13, 14, 15].

Our models’ parameterization focuses on the *Anopheles* species of mosquitoes. These species are noted as the primary vector that transmit the malaria parasite in the African region. The data for temperature-dependent mortality rates in the adult stage originated from a laboratory study to investigate the development and survival of *An. gambiae* at various temperatures [13]. This study monitored mortality rates across a temperature range under various humidity conditions. For our model, we specifically employed the adult mortality data recorded at 60% and 80% humidity, both of which are considered suitable humidity levels for *An. gambiae* mosquitoes [12]. The Gaussian function that best fits the adult mortality data resembles that of the juvenile stage but it is raised to power four instead of power two. This functional form was also used by Beck-Johnson et al, (2013) [12] to model adult mortality. The various functional forms used in describing the life history parameters are given in the main manuscript (see equations 2.4–2.11) with their figures shown in figure 10.

The data used to parameterize the temperature-dependent adult gonotrophic cycle rate were obtained from a research study conducted on *An. pseudopunctipennis* at various constant temperatures [16]. This study consisted of repeated experiments, with each experiment involving 70 to 200 females. These females were force-mated and blood-fed on rabbits, then kept in climatic chambers in individual oviposition vials of approximately 100 ml with cotton soaked in water and covered with filter paper to provide a favorable environment for mosquito egg-laying. The climatic chambers maintained constant temperatures ranging from 15 to 37°C and 70% relative humidity. The experiment was repeated 1 to 3 times. Oviposition vials were checked hourly for the presence of eggs and dead mosquitoes. Individual time to oviposition and mortality were recorded throughout the entire experiment until the last act of egg-laying. Data recorded from this experiment were used to produce parameter estimates (see the function used to represent the data in the main manuscript 2.6 and Figure 10a).

The data for parameterizing temperature- and rainfall-dependent developmental rates were obtained from laboratory studies on *A. gambiae, A. arabiensis*, and *A. pseudopunctipennis*, as compiled by Depinay et al. [17], Parham et al. [10], Bayoh and Lindsay [14], and Okogun et al. [18].

Strains of *A. gambiae s*.*s*., originally from Lagos, were maintained at 26°C and 80% relative humidity. Immature stages were reared at constant temperatures ranging from 10 to 40°C in programmable growth chambers. Water temperature was monitored using data loggers. Adult female mosquitoes were fed on horse blood to produce eggs. Eggs were kept in an environmental chamber at constant temperatures. Hatched larvae were counted and removed daily until no further instars were seen. This procedure was repeated four times for each set temperature. Larvae and pupae were monitored, and development times recorded [14].

Additionally, in an experiment carried out by Okogun et al. [18], mosquito cultures were set up in a pilot study using four different containers: clay pots, metal cans, plastic wares, and bamboo stumps. Detailed descriptions of the experiment are presented in [18]. Environmental temperature, rainfall, and relative humidity were monitored during the study. The female adult mosquitoes were membrane-fed. After two weeks, the number of adult mosquitoes increased, and a baseline count of eggs and larvae was made. Eggs laid were subsequently counted every 2–3 days and transferred to experimental containers kept in netted emergence cages to enable effective monitoring of the developmental stages, through the aquatic stages to the adult stages, and to prevent entry of other mosquitoes. Data recorded from this study were used, in conjunction with the similar experiment by Parham et al. [10], to find parameter estimates for the stage-specific development rates represented in equation 2.8 with their figures shown in figure 10c, d, e.

#### A.2.1 Temperature and rainfall

We assume that temperature and rainfall are seasonal and described by

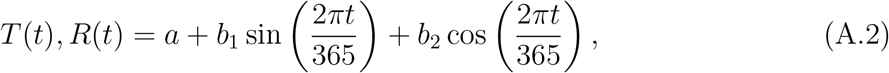

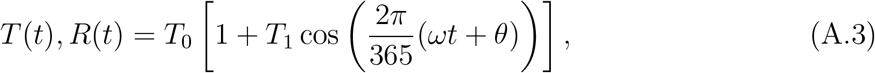

where *a* represent the mean ambient temperature near the surface of the water in degrees Celsius (^*°*^C). It also represent the mean of the cummulative rainfall in mm; *b*_1_ and *b*_2_ affect the magnitude and timing of annual temperature or cummulative rainfall in equation A.2. In equation A.3, *T*_0_ is the mean annual temperature or cummulative rainfall, *T*_1_ captures variations about the mean, *w* and *θ* represent the periodicity and phase shift of the function respectively [1, 4, 8, 19]. For each region, we determine the performance of the fitting by calculating the *R*^2^ values and these are recorded in table 3. Temperature and rainfall data as shown in figure 2 are also recorded in table 4.

**Table 4:**
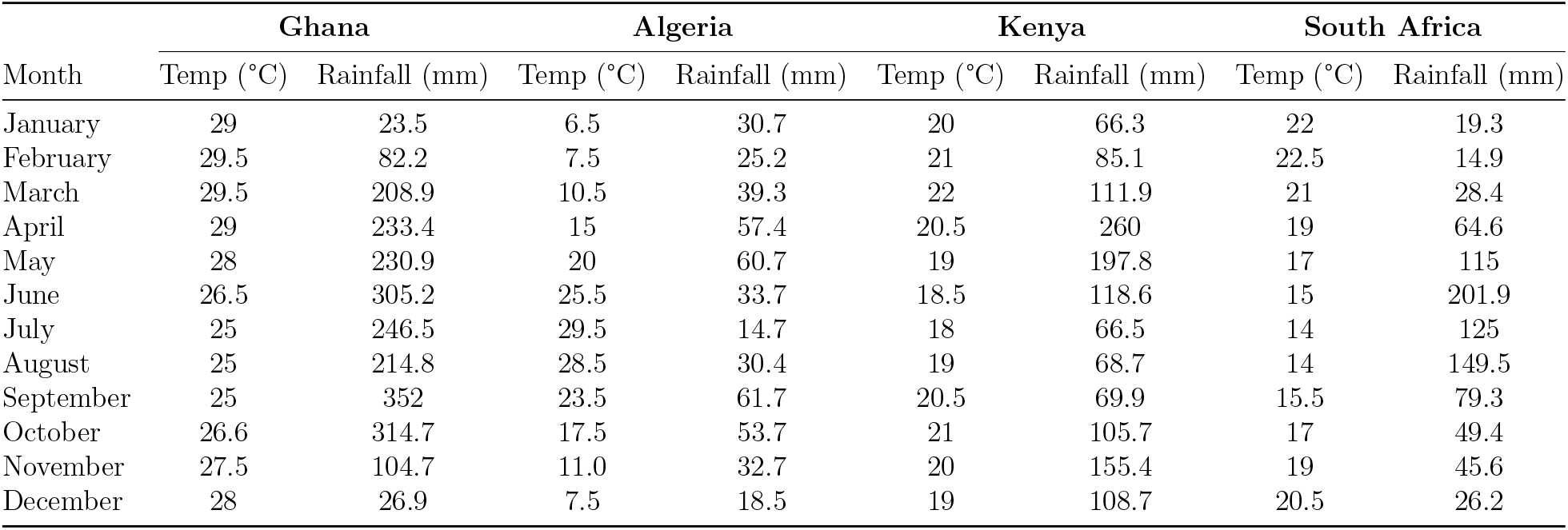
Monthly Average Temperature and Rainfall in Four African Countries. Data is taken from World Weather Online. Sources to specific regions include [20], [21], [22], and [23].

| Month | Ghana |  | Algeria |  | Kenya |  | South Africa |  |
| --- | --- | --- | --- | --- | --- | --- | --- | --- |
|  | Temp (°C) | Rainfall (mm) | Temp (°C) | Rainfall (mm) | Temp (°C) | Rainfall (mm) | Temp (°C) | Rainfall (mm) |
| January | 29 | 23.5 | 6.5 | 30.7 | 20 | 66.3 | 22 | 19.3 |
| February | 29.5 | 82.2 | 7.5 | 25.2 | 21 | 85.1 | 22.5 | 14.9 |
| March | 29.5 | 208.9 | 10.5 | 39.3 | 22 | 111.9 | 21 | 28.4 |
| April | 29 | 233.4 | 15 | 57.4 | 20.5 | 260 | 19 | 64.6 |
| May | 28 | 230.9 | 20 | 60.7 | 19 | 197.8 | 17 | 115 |
| June | 26.5 | 305.2 | 25.5 | 33.7 | 18.5 | 118.6 | 15 | 201.9 |
| July | 25 | 246.5 | 29.5 | 14.7 | 18 | 66.5 | 14 | 125 |
| August | 25 | 214.8 | 28.5 | 30.4 | 19 | 68.7 | 14 | 149.5 |
| September | 25 | 352 | 23.5 | 61.7 | 20.5 | 69.9 | 15.5 | 79.3 |
| October | 26.6 | 314.7 | 17.5 | 53.7 | 21 | 105.7 | 17 | 49.4 |
| November | 27.5 | 104.7 | 11.0 | 32.7 | 20 | 155.4 | 19 | 45.6 |
| December | 28 | 26.9 | 7.5 | 18.5 | 19 | 108.7 | 20.5 | 26.2 |

### A.3 Numerical solution

The system of equations 2.1 were solved using the *deSolve* package in R. We use the ggplot2 package to produce visualization of the model outcomes. Parameter values used in solving the model is presented in table 5.

**Table 5:** Values and functions of the parameters of the model 2.1.

| Parameter | Range/Function Form | References |
| --- | --- | --- |
| $b$ | [150, 300] | [1, 24] |
| $\delta_L$ | [0.02, 0.06] | [4] |
| $\rho_{Ah}$ | [0.6, 2.3] | [2, 25] |
| $\rho_{Ar}$ | [0.5, 1.5] | [2, 25] |
| $K$ | [500, $3 \times 10^8$] | [4, 11, 26] |
| $\mu_{Ao}$ | [0.20, 0.45] | [27] |
| $\mu_{Ah}$ | [0.12, 0.23] | [27] |
| $\phi_{Ao}(T)$ | $\alpha_G \exp \left( -\frac{(T(t)-T_G)^2}{\gamma_G} \right)$ | [12] |
| $\mu_E(T)$ | $\alpha_E \exp \left( \left( \frac{T(t)-T_E}{\gamma_E} \right)^2 \right)$ | [10, 12] |
| $\mu_L(T)$ | $\alpha_L \exp \left( \left( \frac{T(t)-T_L}{\gamma_L} \right)^2 \right)$ | [10, 12] |
| $\mu_P(T)$ | $\alpha_P \exp \left( \left( \frac{T(t)-T_P}{\gamma_P} \right)^2 \right)$ | [10, 12] |
| $\mu_A(T)$ | $\alpha_A \exp \left( \left( \frac{T(t)-T_A}{\gamma_A} \right)^4 \right)$ | [4, 10, 28] |
| $\rho_E(T, R)$ | $\frac{p_E(T, R)}{\tau_E(T)}$ | [1, 4, 10, 18] |
| $\rho_L(T, R)$ | $\frac{p_L(T, R)}{\tau_L(T)}$ | [1, 4, 10, 18] |
| $\rho_P(T, R)$ | $\frac{p_P(T, R)}{\tau_P(T)}$ | [1, 4, 10, 18] |

